# Effects of modified proprioceptive feedback on intra-muscle synergies: An exploration using muscle vibration

**DOI:** 10.64898/2026.09.24.754249

**Authors:** Sayan Deep De, Shanaya Jahanian, Oindrila Sinha, Mark L. Latash

**Author notes:** **Corresponding author:** Sayan Deep De, Department of Kinesiology Rec.Hall-268N, The Pennsylvania State University University Park, PA 16802, USA.

## Abstract

We explored the role of proprioceptive feedback from Ia afferents in intra-muscle synergies stabilizing muscle force with the help of muscle vibration, which is known to increase presynaptic inhibition of those afferents to alpha-motoneurons. Healthy young adults performed an accurate three-finger force production task by pressing with the index, middle and ring fingers of the dominant (right) hand. This was done without vibration and with vibration applied to the extrinsic finger flexors, extensors, both muscles, the palm, or biceps brachii. High-density electromyography was used to identify motor units in both finger flexors and extensors. The raster plots of motor units were converted into firing frequency functions and then subjected to principal component analysis to extract stable motor unit groups (MU-modes). The uncontrolled manifold (UCM) analysis was used to quantify the variance components affecting and not affecting force, and the synergy index in the MU-mode space. Contrary to our expectations, the synergy index did not drop under any vibration condition and showed an increase during vibration applied simultaneously to both flexor and extensor muscles. This was primarily associated with a drop in the component of variance affecting the force magnitude. The results speak against an important role of monosynaptic Ia-afferent projections to alpha-motoneurons in the formation of intra-muscle synergies. They suggest that muscle vibration can play the role of a noisy signal decreasing performance-affecting variance via stochastic resonance effects.

**Highlights:**

- Muscle force is stabilized by covaried recruitment of motor unit groups;
- Indices of force-stabilizing synergies increase under muscle vibration;
- This is due to a drop in the variance component affecting force;
- Intra-muscle synergies are not based on monosynaptic projections of Ia afferents;
- Muscle vibration can improve force accuracy via stochastic resonance.

## Introduction

In his classical book, Bernstein [1] introduced two functions of motor synergies. First, they unite elemental variables (those produced by elements at the selected level of analysis) into robust groups to alleviate the problem of motor redundancy. Such groups have been addressed using different terms such as primitives, factors, modules, modes, etc. ([2–5]; reviewed in[6]). Second, the central nervous system (CNS) acts in the space of modes to organize dynamical stability of salient performance variables, which is crucial for the success of everyday movements given the imperfectly predictable changes in external forces and intrinsic body states. This function of synergies has been explored using a toolbox associated with the uncontrolled manifold (UCM) hypothesis [7]. The most commonly used method compares inter-trial variance in the space of modes that keeps a salient performance variable unchanged (V_UCM_, variance within the UCM) and that leads to changes in this variable (V_ORT_, variance orthogonal to the UCM space). If V_UCM_ > V_ORT_ (both quantified per dimension), a conclusion is drawn about a synergy stabilizing the performance variable. To quantify synergies, an index ΔV = (V_UCM_ – V_ORT_)/(V_UCM_ + V_ORT_) has been commonly used.

Most studies quantified synergies in spaces of kinematic, kinetic, and electromyographic (EMG) variables (reviewed in [8]). Recently, this approach has been generalized to spaces of motor unit (MU) firing frequencies ([9]; reviewed in [10]). Those studies have documented grouping of MUs into MU-modes with parallel scaling of the firing frequencies of individual MUs (confirmed also across muscles, [11,12]) and co-varied recruitment of MU-modes across task repetitions stabilizing muscle force in isometric conditions, confirmed by the synergistic signature V_UCM_ > V_ORT_. Such intra-muscle synergies (IM-synergies) have shown features distinguishing them from the more traditional multi-effector synergies ([13–19]). In particular, IM-synergies stabilize reflex-induced muscle force changes, persist during performance drifts, show no effects of hand dominance, and are reduced under fatigue. In contrast, multi-effector (in particular, multi-finger) synergies do not stabilize reflex-induced muscle force changes, disappear during performance drifts, show effects of hand dominance, and are strengthened under fatigue of an element.

These observations have led to a hypothesis on the existence of distributed parallel circuits stabilizing performance (see also [20]). In particular, multi-effector synergies have been linked to subcortical loops involving the cerebellum and basal ganglia (also supported by studies of neurological patients, reviewed in [21,22]), while IM synergies have been associated with spinal mechanisms such as reflex feedback loops and intraspinal circuits including recurrent inhibition (see also [23,24]). The purpose of our study was to explore the potential role of feedback circuits from peripheral sensory endings sensitive to muscle vibration in the formation of IM synergies.

High-frequency, low-amplitude muscle vibration is known to be a powerful stimulus for several peripheral sensory endings, including primary spindle endings, cutaneous and subcutaneous endings, and Golgi tendon organs [25,26]. It leads to a variety of effects including tonic vibration reflex, stepping-like movements, deviations of the vertical posture, and kinesthetic illusions [27–29]. It has been shown to cause increased presynaptic inhibition of Ia afferent monosynaptic inputs into alpha-motoneurons [30,31]. If those pathways play an important role in IM synergies, indices of these synergies are expected to drop under vibration of the involved muscles (our primary hypothesis).

An earlier study explored effects of vibration on multi-finger synergies during constant total force production tasks by several fingers pressing in parallel in isometric conditions [32]. That study showed a drop in the synergy index ΔV, in particular under vibration applied to the hand. Another study explored effects of vibration on errors in force matching tasks performed by the two hands [33]. It showed qualitatively and quantitatively similar effects during vibration applied to the flexor and extensor muscle groups. This result has suggested that vibration may play the role of a non-specific peripheral noisy input into the CNS. In such a case, the strongest effects can be observed with vibration applied to several of the involved muscles simultaneously (our secondary hypothesis). We tested both hypotheses using a task that required the production of cyclical accurate total force (F_TOT_) by the three fingers pressing in parallel on individual force sensors. The task was performed first without any muscle vibration, then with vibration applied to the hand, extrinsic finger flexors, extensors, both muscle groups simultaneously, and to the biceps of the same arm (a control condition), and finally again without vibration. We expected the IM synergies in the flexor (extensor) to be suppressed strongly by vibration of the flexor (extensor) muscle group.

## Methods

### Subjects

Fourteen healthy individuals (7 females and 7 males; age: 22 ± 2.5 years; mean body mass: 76 ± 16.9 kg; mean height: 1.7 ± 0.09 m) volunteered to participate in this study. Participants were recruited between 12 November 2025 and 20 February 2026. The sample size was selected based on earlier studies examining intra-muscle synergies using the UCM framework, which reported medium-to-large effect sizes, indicating that recruiting 12 participants would provide adequate power [9,19,34]. We recruited additional subjects to account for potential data loss due to the small numbers of motor units identified in the antagonist muscle (extensor digitorum communis, EDC) and possible artifacts from the vibration procedure. All participants were right-handed, as indicated by their preferred hand use for writing and eating with a spoon. Participants had normal or corrected-to-normal vision and reported no history of neurological or musculoskeletal injury affecting the upper extremity. The research was conducted under a protocol approved by the Institutional Review Board (Office for Research Protections, protocol PRAMS00033393) at The Pennsylvania State University, in accordance with the Declaration of Helsinki. All participants provided written informed consent before participation.

### Equipment

Participants were seated in a sturdy chair with back support, their forearms resting comfortably on wooden platforms attached to a rigid table positioned in front of them (see Fig. 1). Three force/torque transducers (Nano-17 sensors, ATI Industrial Automation, Garner, NC, USA) were arranged within a rigid frame (140 × 90 mm) to record the forces of the index (I), middle (M), and ring (R) fingers of the right hand. To prevent slipping, the top surface of each sensor was coated with 320-grit sandpaper. The sensors were spaced 3 cm apart in the mediolateral direction, and adjustments were made in the anterior-posterior direction to align with the individual participant’s finger anatomy. Participants maintained fingertip contact with each sensor with slightly flexed interphalangeal and metacarpophalangeal joints, resulting in a dome-shaped palm. Data acquisition and visual feedback were facilitated through customized software based on the LabVIEW package (LabVIEW version 2014; National Instruments, Austin, TX, USA). Signals from the force sensors were collected, amplified, and sampled at 500 Hz using a 16-bit card (PCI-6225, National Instruments, Austin, TX). Visual feedback was displayed on a 19” monitor positioned 0.6 m in front of the participant at eye level.

**Figure 1.**
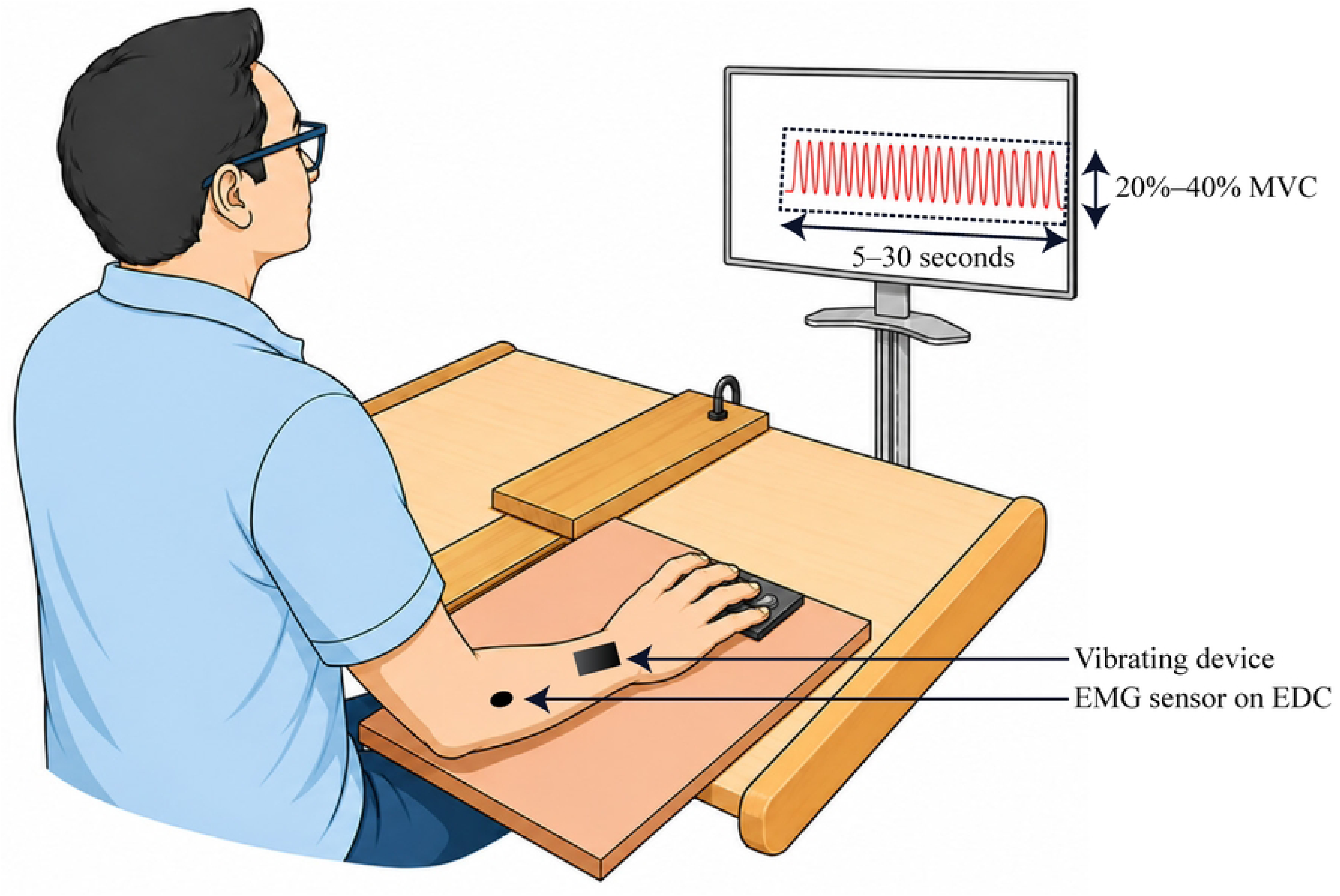
A schematic illustration of the setup with three fingers (index, middle, and ring) pressing on force sensors, the feedback monitor, an approximate location for the electrode on the extensor digitorum communis (EDC) and an approximate location of a vibration over the EDC.

Manual palpation was used to locate the muscle bellies of the flexor digitorum superficialis (FDS) and EDC. The FDS muscle belly was identified between 3 and 7 cm distal to the medial epicondyle along a line towards the lunate, and the EDC muscle belly was identified along a line from the lateral epicondyle to the scaphoid. These regions were marked and prepared by removing hair and cleaning with alcohol swabs to eliminate dead epithelial tissue. Two wireless surface EMG sensors (Trigno Galileo sensor, Delsys Inc., Natick, MA) were attached using removable adhesive stickers over the FDS and EDC muscle bellies. Each sensor body, which houses two stabilizing reference electrodes, was affixed to the distal antebrachium over the distal radius and ulna. The sensor head houses four active electrodes spaced 5 mm apart in a diamond formation, placed over the respective muscle belly. Before testing, EMG electrode location was validated using isometric proximal interphalangeal joint flexion (FDS) and second/third finger metacarpophalangeal extension (EDC) while EMG amplitude was monitored. Isometric distal interphalangeal joint flexion and abduction/adduction of the wrist were additionally used to confirm EMG quiescence of nearby muscles. All EMG signals were initially sampled at 2222.2 Hz with onboard 20–450 Hz filtering. Signal quality was assessed using EMGWorks (Delsys Inc.) and monitored throughout all experimental procedures.

Two electromechanical vibrators (Dynatronic, Marseille, France), each enclosed in a plastic housing accommodating a DC motor with an eccentric on the shaft, were used to deliver mechanical vibration at 100 Hz. Each vibrator weighed 100 g and measured 6 cm in length and 3 cm in diameter. A control unit was used to set the vibration frequency. The vibrators were affixed to the skin over the target locations using elastic straps to ensure consistent contact and prevent slipping throughout each trial. In conditions involving vibration of a single site — the FDS belly, the EDC belly, the palm, or the biceps — only one vibrator was active. In the combined condition, both vibrators were activated simultaneously, one secured over the FDS belly and one over the EDC belly. Placement details per condition are described in the Procedures section. No vibration-induced artifact filtering was applied to the EMG signals. A low-pass filter (50 Hz) was applied to the force signal for real-time visual feedback display during the hand vibration condition only; this filtering was applied exclusively to the on-screen cursor and did not affect the raw data stored for subsequent analysis.

### Procedure

#### Maximum voluntary contraction task

The experiment began with three trials of the maximum voluntary contraction (MVC) task. Visual feedback on the total pressing force (F_TOT_) produced by the index, middle, and ring fingers was provided in the form of a vertical bar on the screen. Participants were instructed to gradually increase their pressing force following an auditory “go” signal, reach their maximal F_TOT_ within 3 s, and sustain this maximum for at least 1 s. A rest period of at least 30 s was provided between trials. The little finger was folded and not used throughout the experiment. The highest F_TOT_ achieved across the three MVC trials was used to set the force targets for the main task.

#### Main task

Participants performed a cyclical three-finger (index, middle, and ring) isometric force production task in which they modulated F_TOT_ between 20% and 40% of their MVC following a sinusoidal target displayed on the screen. Each trial was 30 s in duration, during which participants produced 25 cycles of sinusoidal force at 1 Hz following a 5-second rest period at the start of the trial, during which the fingers remained relaxed on the sensors. Real-time visual feedback on F_TOT_ was provided as a black cursor moving in synchrony with the sinusoidal target. Two trials were collected per condition.

Seven conditions were tested: two no-vibration control blocks administered before and after the vibration conditions (V_N-PRE_ and V_N-POST_, respectively), and five vibration blocks in which the vibrator(s) were applied to the palm (V_H_), FDS (V_F_), EDC (V_E_), both FDS and EDC simultaneously (V_EF_), and the biceps (V_B_). V_N-PRE_ was always administered first and V_N-POST_ always last; the five vibration conditions (V_H_, V_F_, V_E_, V_EF_, V_B_) were presented in a randomized order across participants. Each condition consisted of two trials. For V_H_, V_F_, V_E_, and V_B_, one vibrator was active per trial; for V_EF_, both vibrators were active simultaneously. In all vibration conditions, the vibrator(s) were turned on 30 s prior to trial initiation and remained active throughout the entire recording.

Rest periods of at least 30 s were provided between trials, with longer breaks offered between conditions as needed. The total session duration was approximately 1 hour. No participants reported fatigue.

### Data Analysis

All data were processed using custom-written MATLAB scripts. The analysis involved three sequential steps: (1) extraction of MU action potentials, estimation of the time profile of MU firing frequency, and definition of MU-modes; (2) estimation of Jacobian matrices in both the MU-mode and finger-force spaces; and (3) quantification of force-stabilizing synergies in both spaces using the UCM framework. Each step is described in the following subsections. All force data were digitally low-pass filtered at 10 Hz using a fourth-order, zero-lag Butterworth filter prior to analysis.

### Definition of MU-modes

The EMG signals from the FDS and EDC muscles were decomposed into individual MU spike trains, and the time profiles of MU firing frequencies were estimated using NeuroMap software (Delsys Inc., Natick, MA). Only MUs with accuracy estimates of 80% or higher were retained, as determined by the decompose-synthesize-decompose-compare method implemented within the software. This criterion was adopted for consistency with earlier studies using the same approach. MU decomposition was performed separately for each trial and each muscle, combining the signals collected across all four electrodes of each Trigno Galileo sensor. Trials were excluded from further analysis (17 trials from a total of 196 across all subjects) if fewer than 4 MUs were identified in each muscle.

The discrete MU spike trains were converted to continuous MU firing frequency signals (ƒ_MU_) using a Hann filter with a 1-s window. This smoothing approach enables a continuous representation of MU activity necessary for synergy analysis by integrating the effects of both intermittent MU recruitment and firing rate modulation into a single smooth function. To ensure that the analysis included continuously active MUs, MUs whose firing-rate function, estimated using a 1-s Hann window, fell below 1 Hz at any point during the predefined analysis interval (16%) were excluded from further analysis. Trials were excluded if fewer than two MUs remained in either muscle after this filtering, resulting in the exclusion of an additional 25 trials. Thus, 158 of the 196 planned trial datasets (80.6%) entered the MU-mode analysis.

Force cycles were defined as intervals between two consecutive F_TOT_ peaks. Cycles with durations deviating by more than 20% from 1 s were excluded (8.97%). Note that cycle exclusion did not affect the number of accepted trials. Each accepted cycle was resampled into 20 time bins of equal duration, each representing 5% of the cycle duration. The F_TOT_, individual finger force, and ƒ_MU_ data were averaged within each time bin and then averaged across all accepted cycles within a trial. This yielded an averaged ƒ_MU_ matrix with dimensions of 20 × n, where n is the number of accepted MUs, as well as a 20 × 3 matrix of averaged finger forces.

The analysis of ƒ_MU_ data was performed separately for three MU sets: FDS, EDC, and MUs combined across both muscles (FDS+EDC). For each set, principal component analysis (PCA) was applied to the averaged ƒ_MU_ matrix. Two eigenvectors were retained per trial and per MU set, provided that the following criteria were both satisfied: at least one MU exhibited a loading factor |LF| > 0.5, and the corresponding eigenvalue exceeded 1.0. These criteria were consistently met for two MU-modes across participants and MU sets. For the FDS+EDC combined analysis, one mode exhibited loading factors of opposite sign across MUs from FDS and EDC (reciprocal MU-mode, MU_R_-mode), while the other exhibited same-sign loading factors across both muscle groups (coactivation MU-mode, MU_C_-mode), consistent with prior observations [16,35]. This pattern was observed in at least one trial for each subject. The magnitudes of the two MU-modes for each time bin were obtained by multiplying the ƒ_MU_ matrix by the loading factor vectors of the retained components. Variance accounted for (VAF) by the two MU-modes was computed as a measure of the adequacy of the two-component solution.

### Estimation of the Jacobian matrix

Jacobian in the MU-mode space. The Jacobian matrix (**J**_MU_) represents the partial derivatives of F_TOT_ with respect to changes in MU-mode magnitudes. For each trial and each MU set, multiple linear regression was used to relate changes in F_TOT_ (ΔF) across consecutive 5%-time bins to changes in the two MU-mode magnitudes (ΔM_1_ and ΔM_2_): ΔF = k_1_·ΔM_1_ + k_2_·ΔM_2_, where k_1_ and k_2_ are regression coefficients. The Jacobian matrix was defined as **J**_MU_ = [k_1_ k_2_]. As the MU-mode space was consistently two-dimensional, both the UCM and its orthogonal complement (ORT) were one-dimensional. The null space of **J**_MU_ was used as a linear approximation of the UCM. The coefficient of determination (R²) served as an index of the quality of the Jacobian estimation. The R² > 0.50 inclusion criterion was applied separately to each MU set. Of the 158 trials entering Jacobian estimation, 155 were retained for FDS, 146 for EDC, and all 158 for FDS+EDC. For each participant and condition, the trial with the higher Jacobian VAF was selected for further analysis.

The Jacobian in the finger-force space was computed from trivial mechanics. In the finger-force space, F_TOT_ is the sum of forces produced by the three fingers. Therefore, the Jacobian matrix was defined as **J**_F_ = [1 1 1]. As the finger-force space was three-dimensional, the UCM was two-dimensional, and the ORT was one-dimensional.

### UCM-based analysis in the MU-mode space

The UCM hypothesis framework [7] was used to quantify F_TOT_-stabilizing synergies in the MU-mode space. For each trial, mean-free vectors in MU-mode space were obtained by subtracting the average values across cycles for each 5% time bin. These vectors were then projected onto the UCM (the null space of **J**_MU_) and onto the ORT, yielding inter-cycle variance components V_UCM_ and V_ORT_, respectively. V_UCM_ reflects variance in MU-mode space that leaves F_TOT_ unchanged, while V_ORT_ reflects variance that produces changes in F_TOT_. No normalization per dimension was required, as both UCM and ORT were one-dimensional. The UCM-based analysis was performed separately for MU-modes derived from FDS, EDC, and FDS+EDC.

### UCM-based analysis in the finger-force space

A similar UCM-based analysis was performed in the space of individual finger forces. Mean-free finger-force vectors were projected onto the UCM (the two-dimensional null-space of **J**_F_) and onto the one-dimensional ORT, yielding V_UCM_ and V_ORT_ in the finger-force space. Both variance components were normalized per dimension prior to the computation of the synergy index to account for the difference in dimensionality between the UCM and ORT subspaces.

### Outcome variables

For each space, the synergy index was defined as: ΔV = (V_UCM_ − V_ORT_)/(V_UCM_ + V_ORT_). Since ΔV is bounded between −1 and +1, Fisher’s z-transformation was applied prior to statistical analyses: ΔV_Z_ = 0.5 × ln[(1 + ΔV)/(1 − ΔV)]. This transformation yields ΔV_Z_ = 0 in the absence of a synergy. A positive ΔV_Z_ indicates a force-stabilizing synergy (V_UCM_ > V_ORT_). The primary outcome variables were V_UCM_, V_ORT_, and ΔV_Z_, computed separately for the MU-mode space (for each of the three MU sets: FDS, EDC, and FDS+EDC) and for the finger-force space.

### Statistics

Statistical analyses were performed in R using RStudio (2023.06.0+421; Posit Software, PBC). Data were checked for normality and sphericity before parametric analyses. Greenhouse–Geisser corrections were applied when sphericity was violated. V_UCM_ and V_ORT_ were natural-log-transformed before statistical analysis.

One-way repeated-measures ANOVAs were used to examine the effect of Condition on RMSE and the finger-force-space synergy variables. The primary synergy analyses compared V_F_, V_E_, V_EF_, and V_N-POST_. PCA and Jacobian VAF values were analyzed using linear mixed-effects models with Condition, Muscle (FDS, EDC, and FDS+EDC), and their interaction as fixed effects and participant as a random factor. MU-mode-space ΔV_Z_, V_UCM_, and V_ORT_ were analyzed separately for FDS, EDC, and FDS+EDC using mixed-effects models with Condition as a fixed effect and participant as a random factor. For these models, denominator degrees of freedom were estimated using the Satterthwaite approximation.

Significant effects were explored using pairwise comparisons with Holm correction. Statistical significance was accepted at p < 0.05. Partial eta squared (η²_p_) was reported as the effect-size measure. Data are presented as mean ± SE unless otherwise stated.

## Results

### Typical behavior

All participants successfully performed the cyclical three-finger force production task across all conditions, modulating F_TOT_ between 20% and 40% MVC in synchrony with the sinusoidal target. Figure 2 shows an exemplary trial from a representative participant illustrating the force profile relative to the task template.

**Figure 2.**
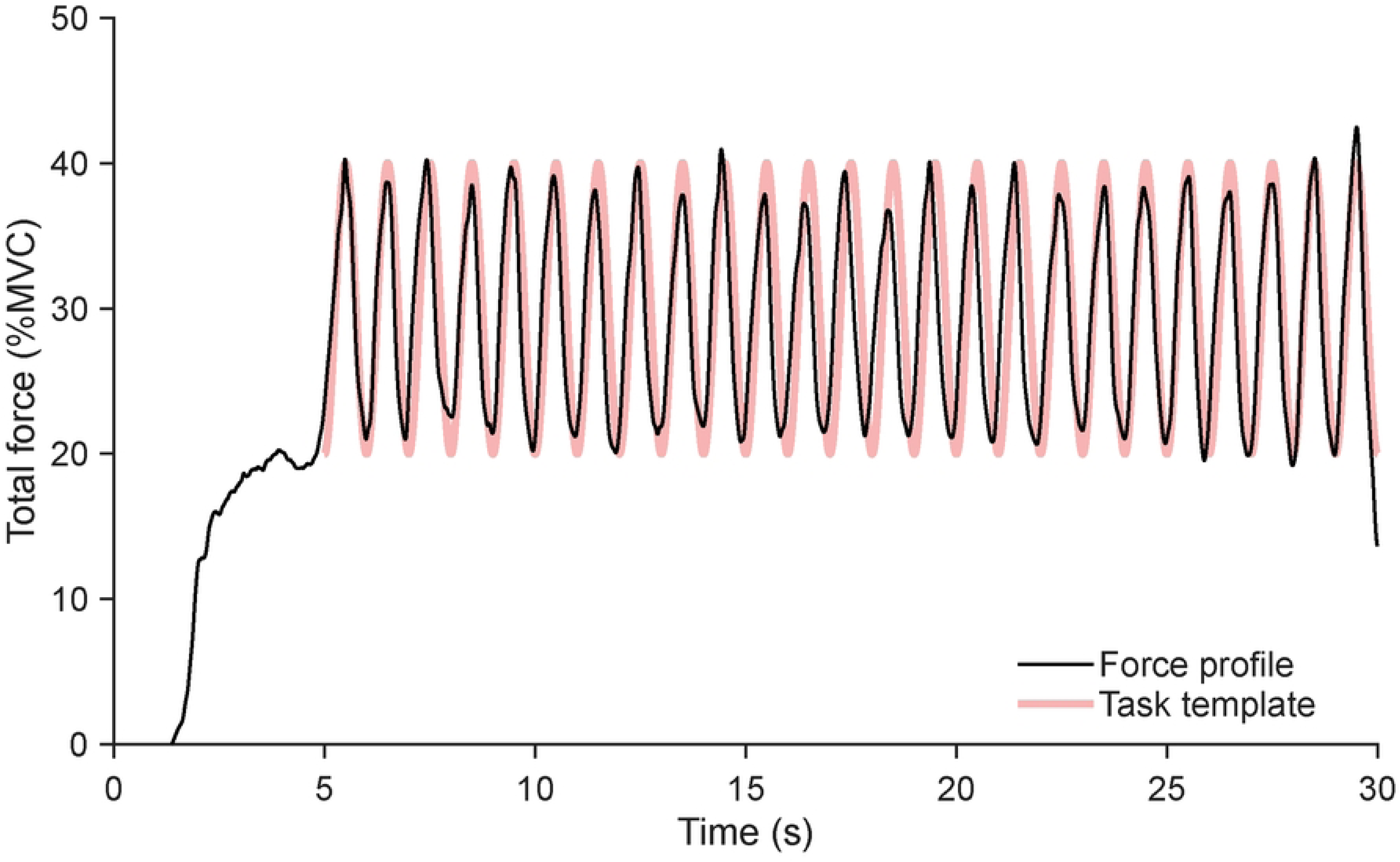
An example of typical performance by a representative subject. A single trial is shown performed without vibration.

Performance accuracy was quantified using RMSE computed relative to both the participant’s own average force profile (RMSE_AVG_) and the task template (RMSE_TEMP_). These results are illustrated in Figures 3A and 3B, respectively. RMSE values relative to the average force profile (RMSE_AVG_) showed a significant effect of Condition, reflecting higher values for the first trial (V_N-PRE_) compared to most other conditions (F_[3.51, 42.15]_ = 6.77, p < 0.001, η²_p_ = 0.361). Post-hoc comparisons revealed that RMSE_AVG_ was higher during V_N-PRE_ (2.39 ± 0.18 %MVC) relative to V_F_ (2.04 ± 0.13 %MVC, p = 0.002), V_E_ (1.99 ± 0.11 %MVC, p < 0.001), V_EF_ (2.00 ± 0.13 %MVC, p < 0.001), V_B_ (2.02 ± 0.13 %MVC, p < 0.001), and V_N-POST_ (1.89 ± 0.11 %MVC, p < 0.001). The difference between V_N-PRE_ and V_H_ (2.17 ± 0.12 %MVC) did not reach significance (p = 0.245). RMSE_AVG_ values for V_H_ were higher than for V_N-POST_ (p < 0.05). No significant differences were found among the other vibration conditions (all p > 0.05).

**Figure 3.**
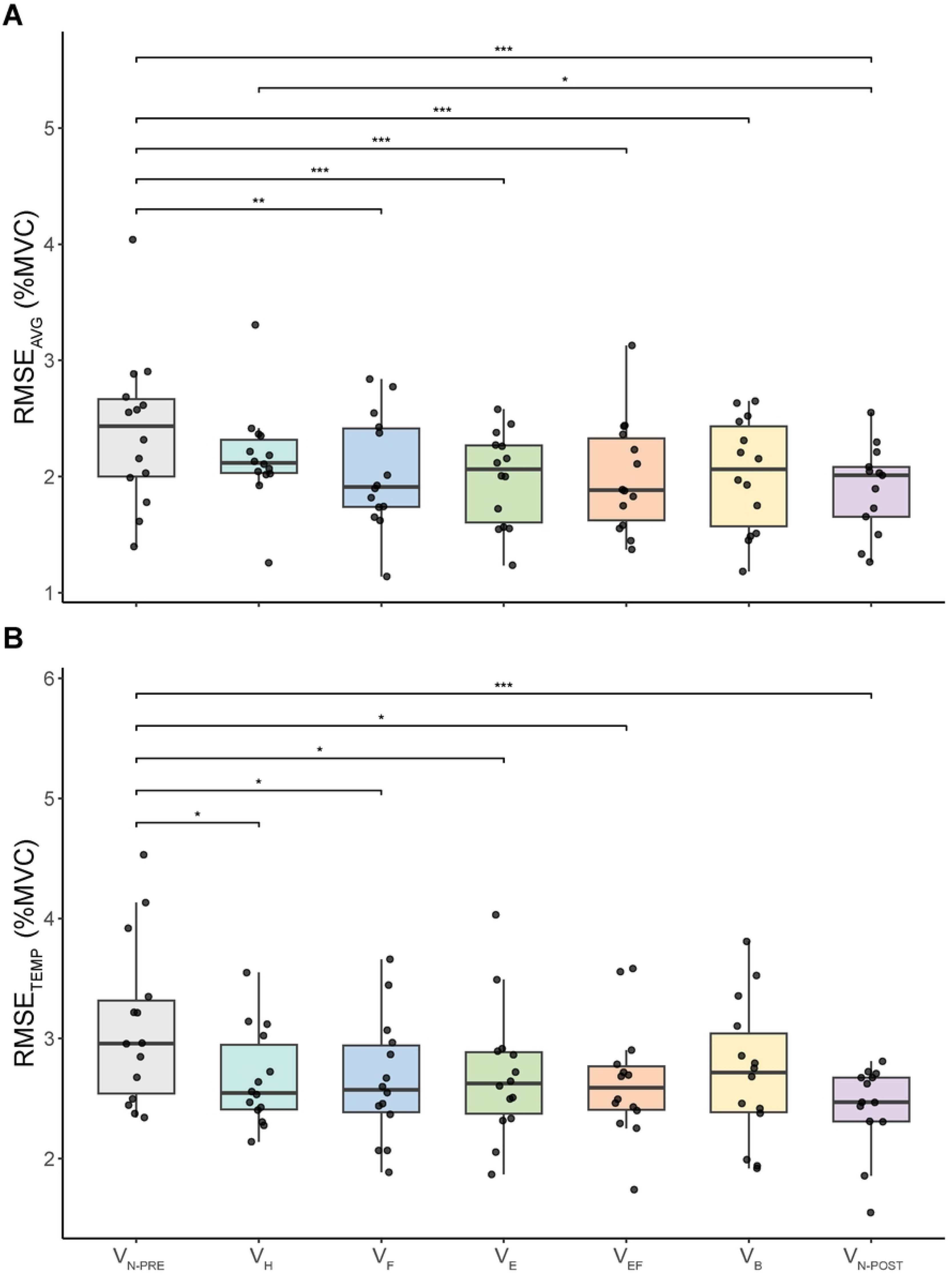
Root mean square error (RMSE) computed with respect to the average cycle performance (panel A) and with respect to the template shown on the screen (panel B). Both plots show both the data for individual subjects (small black dots) and box-with-whiskers values about the medians. * - p < 0.05; ** - p < 0.01; *** - p < 0.001. The X-axis shows the conditions: V_N-PRE_ – the first conditions without vibration, V_H_ - vibration of the hand, V_F_ – vibration of the flexor digitorum superficialis, V_E_ – vibration of the extensor digitorum communis, V_EF_ – vibration of both flexor and extensor muscles, V_B_ – vibration of the biceps brachii, V_N-POST_ – the last condition with no vibration.

A similar pattern was observed for RMSE computed relative to the task template (RMSE_TEMP_), which also showed a significant effect of Condition (F_[2.75, 33.03]_ = 4.91, p = 0.007, η²_p_ = 0.290). RMSE_TEMP_ was higher during V_N-PRE_ (3.10 ± 0.18 %MVC) relative to V_H_ (2.67 ± 0.11 %MVC, p = 0.024), V_F_ (2.65 ± 0.14 %MVC, p = 0.017), V_E_ (2.70 ± 0.15 %MVC, p = 0.050), V_EF_ (2.64 ± 0.13 %MVC, p = 0.014), and V_N-POST_ (2.43 ± 0.36 %MVC, p < 0.001). The difference between V_N-PRE_ and V_B_ (2.71 ± 0.16 %MVC) did not reach significance (p = 0.074). No significant differences were observed among the other vibration conditions (all p > 0.05).

Taken together, these results indicate that tracking error was slightly higher at the start of the session (V_N-PRE_) and then stabilized, with performance under vibration conditions comparable to V_N-POST_ across both accuracy measures. Accordingly, V_N-PRE_ was considered a practice condition. Preliminary analyses showed no differences in the synergy outcome variables involving either of the two control conditions, vibration applied to the palm of the hand (V_H_) or vibration applied to the biceps (V_B_). Therefore, the subsequent presentation of the synergy results focuses on four conditions, V_F_, V_E_, V_EF_, and V_N-POST_.

### Multi-finger synergies

When averaged across the 20 phases of the force cycle, analysis in the finger-force space revealed an absence of a force-stabilizing multi-finger synergy, as reflected by V_UCM_ < V_ORT_ and negative ΔV_Z_ values across V_F_, V_E_, V_EF_, and V_N-POST_ (Figure 4). Mean ΔV_Z_ ranged from −0.211 ± 0.106 in V_F_ to −0.135 ± 0.078 in V_N-POST_ (mean ± SE). The analysis included 11 participants with complete data across the four conditions. A one-way repeated-measures ANOVA showed no effect of Condition on ΔV_Z_ (F_[2.60, 25.99]_ = 0.20, p = 0.873, η²_p_ = 0.019).

**Figure 4.**
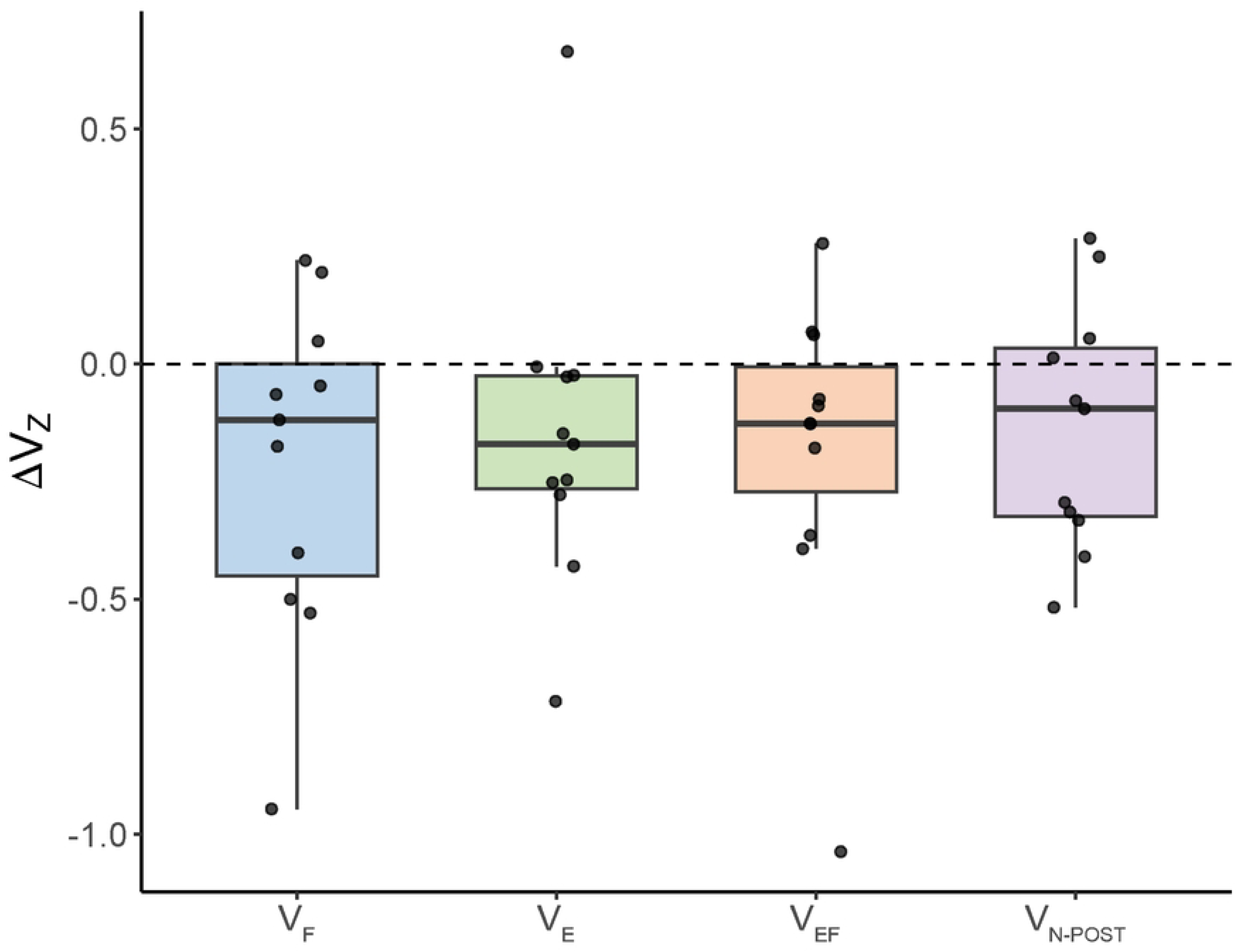
The synergy index (ΔV_Z_) after Fisher’s z-transformation computed within the space of individual finger forces. The data for individual subjects (small black dots) and box-with-whiskers values about the medians are shown. V_F_ – vibration of the flexor digitorum superficialis, V_E_ – vibration of the extensor digitorum communis, V_EF_ – vibration of both flexor and extensor muscles, V_N-POST_ – the last condition with no vibration.

Analysis of the natural-log-transformed variance components, also averaged across the 20 phases of the force cycle, showed no effect of Condition on V_UCM_ (F_[2.40, 23.99]_ = 0.46, p = 0.672, η²_p_ = 0.044) or V_ORT_ (F_[2.20, 21.97]_ = 0.36, p = 0.723, η²_p_ = 0.034).

### MU-mode-based synergies

#### MU-identification and PCA

Across V_F_, V_E_, V_EF_, and V_N-POST_, 87 of the 112 performed trials (77.7%) entered the MU analysis; the remaining 25 trials (22.3%) were unavailable because not enough MUs had been identified or had not passed the frequency-based inclusion criteria. PCA consistently identified two MU-modes when performed separately for FDS, EDC, and the combined FDS+EDC sets of motor units. Averaged across the four conditions, the two modes accounted for 82.6 ± 2.1% of the variance in FDS, 87.6 ± 3.1% in EDC, and 79.6 ± 2.7% in the combined FDS+EDC analysis (mean ± SE; Figure 5A).

**Figure 5.**
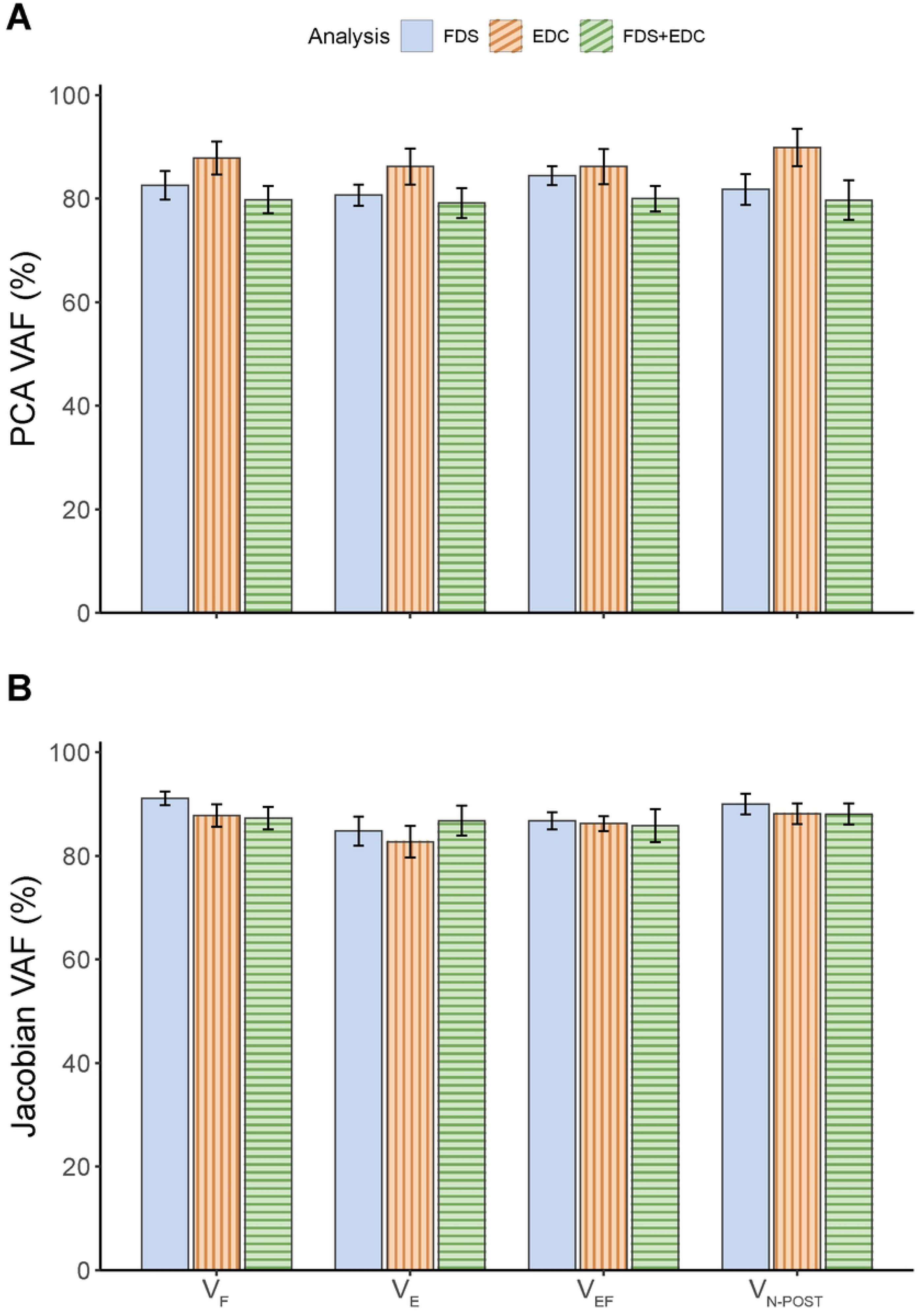
The amount of variance accounted for at the stages of principal component analysis (PCA VAF, panel A) and Jacobian identification (Jacobian VAF, panel B) in the space of motor units within the flexor digitorum superficialis (FDS; blue, solid bars), extensor digitorum communis (EDC; orange, vertically-striped bars), and both muscles together (FDS+EDC; green, horizontally-striped bars). Means ± standard error bars are shown. V_F_ – vibration of the flexor digitorum superficialis, V_E_ – vibration of the extensor digitorum communis, V_EF_ – vibration of both flexor and extensor muscles, V_N-POST_ – the last condition with no vibration.

A two-way linear mixed-effects analysis showed no effect of Condition on PCA VAF (F_[3, 116.14]_ = 0.34, p = 0.794, η²_p_ = 0.009) and no Condition × Muscle interaction (F_[6, 116.04]_ = 0.58, p = 0.746, η²_p_ = 0.029). The effect of Muscle was significant (F_[2, 116.04]_ = 20.57, p < 0.001, η²_p_ = 0.262). Holm-adjusted comparisons showed that PCA VAF was higher for EDC than for FDS (p < 0.001) and FDS+EDC (p < 0.001), and higher for FDS than for FDS+EDC (p = 0.020).

#### Jacobian Estimation

The coefficient of determination (R²) was used to assess the quality of the Jacobian estimation, with R² > 0.50 used as the inclusion criterion. Of the 87 trials entering this stage, 85 (97.7%) were retained for FDS, 83 (95.4%) for EDC, and all 87 (100%) for FDS+EDC. Averaged across the four conditions, Jacobian VAF was 88.1 ± 1.1% for FDS, 85.9 ± 1.4% for EDC, and 86.8 ± 2.0% for FDS+EDC (mean ± SE; Figure 5B). The linear mixed-effects analysis showed no effect of Condition (F_[3, 116.44]_ = 2.22, p = 0.089, η²_p_ = 0.054), no effect of Muscle (F_[2,115.87]_ = 0.91, p = 0.407, η²_p_ = 0.015), and no Condition × Muscle interaction (F_[6, 115.86]_ = 0.45, p = 0.842, η²_p_ = 0.023).

#### Force-stabilizing synergies in the MU-mode space

When averaged across the 20 phases of the force cycle, ΔV_Z_ was positive in V_F_, V_E_, V_EF_, and V_N-POST_ for FDS, EDC, and FDS+EDC, indicating force-stabilizing synergies in the MU-mode space (Figure 6). Depending on the muscle and condition, data from 11 or 12 participants were available for the mixed-effects analyses. No effect of Condition on ΔV_Z_ was found for FDS (F_[3, 31.02]_ = 1.09, p = 0.369, η²_p_ = 0.095). Significant condition effects were found for EDC (F_[3, 31.21]_ = 4.25, p = 0.013, η²_p_ = 0.290) and FDS+EDC (F_[3, 32.23]_ = 3.43, p = 0.029, η²_p_ = 0.242). Holm-adjusted comparisons showed that EDC ΔV_Z_ was higher in V_F_ (p = 0.032) and V_E_ (p = 0.035) than in V_N-POST_. In the FDS+EDC analysis, ΔV_Z_ was higher in V_EF_ than in V_N-POST_ (p = 0.024).

**Figure 6.**
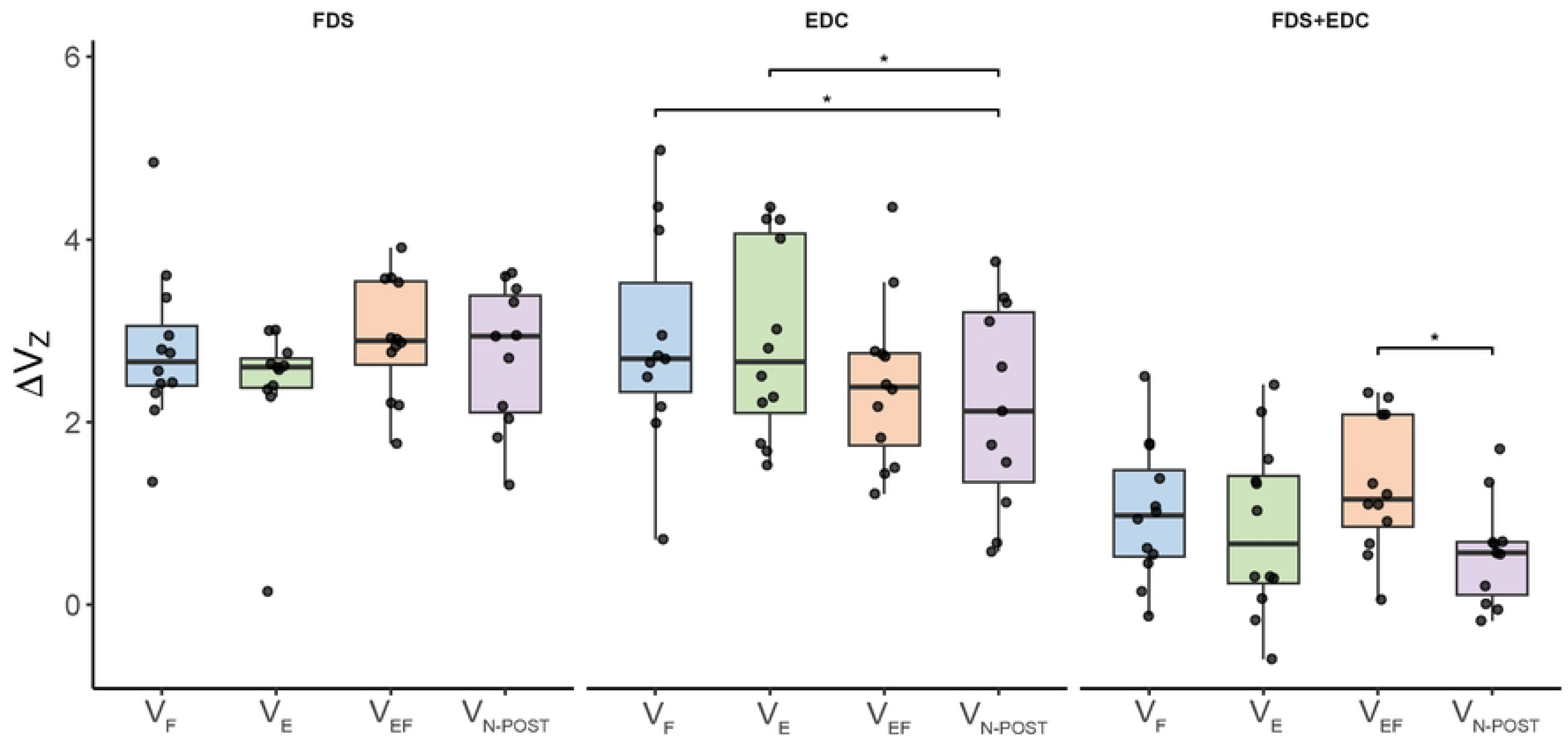
The synergy index (ΔV_Z_) after Fisher’s z-transformation computed within the space of motor unit modes (MU-modes) defined for the flexor digitorum superficialis (FDS), extensor digitorum communis (EDC) and both muscles. The data for individual subjects (small black dots) and box-with-whiskers values about the medians are shown. V_F_ – vibration of the FDS, V_E_ – vibration of the EDC, V_EF_ – vibration of both FDS and EDC, V_N-POST_ – the last condition with no vibration. * – p < 0.05.

Analysis of the natural-log-transformed variance components (Figure 7) showed no effect of Condition on V_UCM_ (Figure 7A) for FDS (F_[3, 31.65]_ = 1.33, p = 0.281, η²_p_ = 0.112), EDC (F_[3, 31.32]_ = 2.80, p = 0.056, η²_p_ = 0.212), or FDS+EDC (F_[3, 32.18]_ = 2.56, p = 0.073, η²_p_ = 0.192). For V_ORT_ (Figure 7B), no condition effect was found for FDS (F_[3, 30.84]_ = 1.00, p = 0.407, η²_p_ = 0.088). Significant condition effects were found for EDC (F_[3, 31.25]_ = 3.86, p = 0.018, η²_p_ = 0.271) and FDS+EDC (F_[3, 32.21]_ = 5.86, p = 0.003, η²_p_ = 0.353). For EDC, V_ORT_ was lower in V_F_ and V_EF_ than in V_N-POST_ (both p = 0.036). For FDS+EDC, V_ORT_ was lower in V_F_ (p = 0.014) and V_EF_ (p = 0.003) than in V_N-POST_. Thus, the higher synergy indices were observed without significant changes in V_UCM_ and were accompanied by lower variance affecting total force.

**Figure 7.**
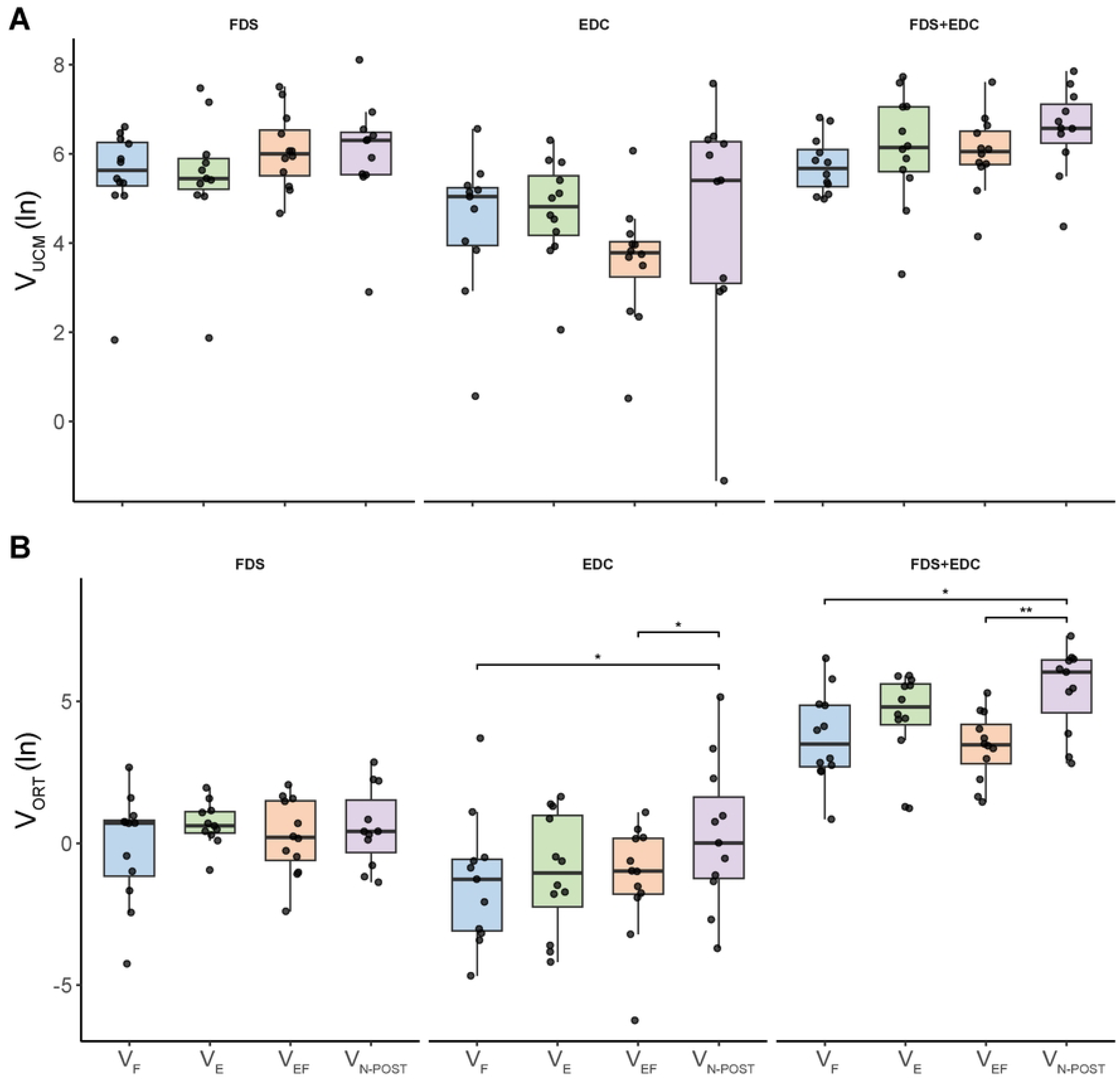
Natural log-transformed values of the two components of inter-cycle variance, along the uncontrolled manifold (panel A, V_UCM_) and orthogonal to the uncontrolled manifold (panel B, V_ORT_) for the MU-modes defined for the flexor digitorum superficialis (FDS), extensor digitorum communis (EDC) and both muscles. The data for individual subjects (small black dots) and box-with-whiskers values about the medians are shown. V_F_ – vibration of the FDS, V_E_ – vibration of the EDC, V_EF_ – vibration of both FDS and EDC, V_N-POST_ – the last condition with no vibration. * – p < 0.05; ** – p < 0.01.

## Discussion

The data clearly disprove our main hypothesis by showing effects of vibration in the direction opposite to the predicted one. Indeed, while the main hypothesis predicted a drop in the MU-mode force-stabilizing synergy index (ΔV) under vibration due to the increased presynaptic inhibition of Ia-afferent input to alpha-motoneurons [30,31], the experiment showed either no statistically significant changes (within the space of MU-modes for FDS) or a significant increase under the vibration applied to either FDS or EDC (within the space of MU-modes for EDC), or to both extrinsic flexor and extensor muscle groups (within the space of MU-modes defined for the combined sets of MUs across the two muscles). Our secondary hypothesis has been confirmed for the space of MU-modes across the two muscles (FDS+EDC): We observed stronger and more consistent effects of vibration on the IM-synergy index when the vibration was applied to the flexor and extensor muscles simultaneously as compared to the effects of vibration to the extrinsic finger flexor alone. We did not observe comparable effects with vibration applied to one of the muscles.

Another unexpected finding has been the predominant role of changes in the variance component affecting performance (V_ORT_) in the observed effects on ΔV. This is in contrast to several studies pointing at differences in the variance component with no effects on performance (V_UCM_) as the main contributor to changes in the synergy index following practice, in neurological patients, and reflecting effects of dopamine-replacement drugs [36–39]. We offer an interpretation of this effect based on the phenomenon of stochastic resonance discussed in a number of studies of the neural control of movements [40,41].

### Potential circuits involved in IM-synergies

Performance-stabilizing synergies can get contributions from both experience-based feed-forward mechanisms and feedback-based loops [42–44]. The feed-forward mechanisms have been shown to lead to variable across trials sharing of task-related performance variables among the contributing elements, seen from the very beginning of performance [45]. The variable sharing contributes to V_UCM_ and persists over relatively long times, even when the salient feedback is removed, leading to drifts in the performance variable [46,47] and disappearance of performance-stabilizing synergies (V_UCM_ ≈ V_ORT_) [48]. The feedback-based effects are reflected in negatively covarying contributions of elements during steady-state task performance. These effects disappear after the removal of the salient sensory feedback [48,49]. So far, these conclusions have been based on studies of multi-effector synergies using manipulations of the visual feedback in tasks involving both upper and lower extremities (see also [33,50]).

There is little evidence pointing at specific circuits involved in IM-synergies. The contrasting findings in studies of multi-effector synergies and IM-synergies have strongly suggested that the IM-synergies are primarily based on spinal circuitry. This conclusion has been supported by the lack of hand dominance effects [16,17] and the effects of presenting or removing the salient visual feedback on the indices of IM-synergies [19]. On the other hand, IM-synergies stabilize reflex-induced force changes, while multi-effector synergies fail to show such effects [17]. One of the results of our study, an increase in the synergy index under the vibration of both extrinsic flexor and extensor muscle groups, is qualitatively different from a drop in the force-stabilizing multi-finger synergy index under muscle vibration reported earlier [32]. This finding can also be seen as corroborating the hypothesis on different neural circuitry involved in IM-synergies and multi-finger synergies.

There are many spinal circuits that can potentially contribute to IM-synergies. These include the negative feedback loops from proprioceptive sensory endings and intra-spinal circuits such as the recurrent inhibition. The latter mechanism has been discussed as a potential contributor to the stability of the alpha-motoneuronal pool output [23].

Our experiment has provided strong evidence against a major role of Ia-afferent monosynaptic inputs into alpha-MNs in IM-synergies. Indeed, the documented increase in presynaptic inhibition of these projections by muscle vibration [30,31] failed to have any negative effects on the synergy index. There can be effects from other reflex-mediated sources such as polysynaptic inputs into alpha-MNs, in particular those involved in the tonic vibration reflex, which may be enhanced during muscle vibration. Possible other sources include reflex effects from Ib afferents, which respond to muscle vibration in the presence of non-zero baseline muscle activation [26] and could be particularly relevant for accurate force production tasks. No inhibitory effects of vibration have been described for those loops. Intraspinal circuitry such as recurrent inhibition, reciprocal inhibition, etc. could play a major role in IM-synergies. Note that the V_EF_ condition could be expected to increase the reciprocal inhibition effects by the increase in the Ia afferent input. The larger effects of V_EF_, as compared to the no-vibration and flexor vibration conditions, on the synergy index are consistent with this interpretation.

### Higher stability under vibration: Potential role of stochastic resonance

Arguably, the most unexpected result of the study is the increase in the IM-synergy index under the vibration applied simultaneously to the extrinsic flexor and extensor muscle groups or to the antagonist muscle (EDC) alone. Indeed, muscle/tendon vibration has been traditionally viewed as a factor leading to kinesthetic errors in coordinate [28,51], velocity [51], and force [52,53] affected by muscles subjected to the vibration. Most cited studies linked the observed effects to the vibration-induced increase in the firing frequency of primary spindle endings. Later, effects of vibration on the efferent component of kinesthetic perception were suggested and received experimental confirmation [33,54,55]. Nevertheless, the consensus has been that muscle vibration is a detrimental factor for any task involving accurate production and/or perception of a mechanical variable to which the muscle group with the vibrator contributes.

There are, however, many observations suggesting that vibration effects are not limited to the most commonly invoked phenomena such as the tonic vibration reflex, increase in the presynaptic inhibition of Ia-afferent monosynaptic projection on alpha-MNs, and perceptual errors in the direction expected from the responses of the sensory endings to vibration. In particular, reflex activation can be seen in muscles not directly affected by vibration [56,57], patterns of muscle activation can be complex, including cyclical leg movements, suggesting involvement of a central pattern generator for locomotion [29,58], and kinesthetic illusions can switch direction in the presence of apparently irrelevant stimuli [55]. A recent study explored errors in matching forces produced by the four fingers of a hand with the fingers of the other hand and reported similar error patterns under vibration applied to the extrinsic flexor or extensor muscle groups [33]. The authors of the study have concluded that the vibration played the role of a noise factor, with the source of the noise being relatively unimportant for the induced perceptual and motor effects.

Can vibration-induced “noise” be helpful in the organization of force-stabilizing IM synergies? In particular, can vibration amplify subtle effects at the spinal level using the stochastic resonance phenomenon? Such phenomena have been explored and used in clinical studies of impaired postural control, which improved when noisy signals were applied to the feet, to the whole body, or to the vestibular apparatus [40,41,59,60]. The idea is that adding noise increases the chances that a small, useful signal reaches the threshold for detection. Note that the increased synergy index in our study was associated with a drop in the component of variance (V_ORT_) that affected force without a comparable change in the component that had no effect on force magnitude (V_UCM_). If the structure of variance is defined by spinal circuitry, effectiveness of this mechanism can be increased under the noise factor induced by the vibration. Note that significant changes in the synergy index were seen under the vibration applied simultaneously to both flexor and extensor muscles when the assumed noise factor could be expected to be maximal.

The increase in the synergy index associated with a drop in V_ORT_ is by itself unexpected. Indeed, many earlier studies documented differences in V_UCM_ under a variety of factors including neurological disorder [39], effects of drug treatment [38], and short-term practice [36], while changes in V_ORT_ were relatively small and inconsistent. Note that the synergy index reflects the relative magnitudes of V_UCM_ and V_ORT_, not their absolute magnitudes, i.e., how successful the system is in channeling the unavoidable variance at the level of elements into the UCM where the variance has no effects on the important performance variable.

### Concluding comments and methodological issues

The exact nature of IM-synergies remains unknown. Our results argue against one of the potential contributors (monosynaptic Ia-afferent inputs) and suggest an unexpected possible contributor – stochastic resonance induced by an overall increased diffuse afferent inflow. Future studies may be able to analyze possible contributions of intra-spinal circuitry and polysynaptic reflex feedback loops.

We must admit a few drawbacks of this study, which are inherent to nearly all studies that use surface electrodes to identify motor units and quantify their firing frequency. First, the process of MU identification is imperfect. We used the setting of 80% confidence, which had been used in a number of earlier studies [9,16,17,35]. This setting is a trade-off between achieving a relatively high level of confidence and the ability to identify a reasonable number of MUs that allows performing the next steps in data processing. The method of estimating the MU instantaneous firing frequency with the help of the Hann filter is also imperfect. It combines the frequency modulation and recruitment into a single mechanism and is associated with a rather poor time resolution. On the other hand, some MUs can fire at frequencies of a few Hz, which by itself limits the time resolution since estimating instantaneous frequency requires observing at least two action potentials within any given time window. A major limitation of the current setup and software is that it is impossible to identify MUs across trials. Hence, MU-modes and Jacobians had to be defined separately across conditions. We saw no major effects of vibration on quantifiable characteristics at the steps of MU-mode identification and Jacobian estimation. In particular, variance accounted for at each of those steps was comparable across the vibration conditions. In spite of all the mentioned drawbacks, we believe that the main significant findings of the study are reliable, although confirmation in future studies is very much desirable.

## Statements and Declarations

### Ethics approval

The study protocol was approved by the Institutional Review Board of the Pennsylvania State University (Protocol PRAMS00033393).

### Consent to participate

Written informed consent was obtained from all participants.

## Generative AI use

During the preparation of this work, one of the authors (S.D.D.) used Claude (Anthropic) and ChatGPT (OpenAI) to improve the wording and language of the manuscript.

After using these tools, all authors reviewed and edited the content as needed and take full responsibility for the content of the published article.

## Author contributions

SDD – study design, data collection, data processing, writing the manuscript, editing the manuscript, approval of the final version;

SJ – data collection, data processing, approval of the final version.

OS – data processing, editing the manuscript, approval of the final version.

MLL – study design, data processing, editing the manuscript, approval of the final version.

## Data availability

All data underlying the findings and the analysis code are publicly available from the Open Science Framework (OSF) at https://doi.org/10.17605/OSF.IO/B823E.

